# Newly developing methodologies to investigate health hazards posed by ionizing radiation to human space missions

**DOI:** 10.1101/2022.03.28.486099

**Authors:** Franco Ferrari, Ewa Szuszkiewicz

## Abstract

Ionizing radiation is one of the main threats to human space exploration beyond low Earth orbit (BLEO). It is thus of primary importance to determine safe career dose limits for astronauts involved in BLEO missions. In the first part of this work it is shown how the methods of physics and statistics can contribute to its solution. The average equivalent doses received by a hypothetical human crew are established using the data of several robotic missions to the Moon and to Mars. The probabilities of the occurrence of deterministic effects due to radiation that could impair the success of a mission or lower the life expectancy of astronauts are evaluated with the help of a statistical analysis. In the last part of this work it is argued that the use of the so-called 3D models or organoids combined with the methods of precision oncology and molecular medicine could be a good candidate of a strategy in order to predict the insurgence of stochastic effects in humans. On one side, organoids recapitulate several features of the real human organs and their in vivo surroundings. On the other side, we argue with a case study that precision oncology and molecular medicine are able to provide a deeper insight of the onset of cancer following irradiation in space.

## Introduction

The risks posed by energetic particle radiation to human crews implementing exploration missions in Low Earth Orbit (LEO) and Beyond Low Earth Orbit (BLEO) are of a serious nature. Both the short-term and long-term effects of irradiation in space should be taken into account when establishing safe career dose limits for astronauts, see [1, 2].

An obstacle in estimating for astronauts the maximum acceptable dose accrued in interplanetary space below which the risk of contracting cancer or other pathologies is not higher than the risk incurred by workers on Earth dealing with sources of ionizing radiation, is the lack of sufficient statistics concerning the effects of high LET radiation on human space travelers. Hazardous radiation (including energetic protons and heavy ions) is present outside the region protected by the geomagnetic field (i.e. outside the Earth ‘s magnetosphere) and, hitherto, data concerning the effects of this ionizing radiation on living organisms in BLEO has not been available - apart from measurements made during the Apollo lunar exploration missions of the 1960s and 1970s, see for example [3]. An experiment that is expected to quantify the level of biological damage incurred inspace, as well as to validate existing models, is conducted aboard NASA ‘s BioSentinel nanosatellite, utilizing a payload launched in November 2022.

In the present work we propose a few ideas from a physicist ‘s point of view in order to tackle the problem of evaluating the health hazards posed by ionizing radiation to human space missions to BLEO objects. The aim is to see how far it is possible to push physical considerations in order to obtain new knowledge on this subject. The obtained results are not meant to substitute the existing and very sophisticated methods for determining the safe career doses of ionizing radiation for astronauts that are already exploited. An example of such methods is provided by the NASA Space Cancer Risk model (NSCR-2022). A few other models will be briefly described below. The interested reader may also consult [4] for a recent review on this topic.

## Methods

To begin with, we provide a rough estimation of the equivalent doses that are expected to be received by a hypothetical crew during the travel to the Moon or to Mars and a short stay on these objects for exploration purposes. In doing this, we do not take into account the large Solar Energetic Particle (SEP) events. Of course, these events may significantly increase the risks for astronauts. On the other side, it is difficult to quantify the effects on space traveling of such irregular phenomena whose occurrences cannot be predicted so far. For the interested reader, a more extensive account of radiation exposure for different scenarios can be found in [5, 6].

In order to compute the typical doses incurring during a mission, it is of critical importance to evaluate not only doses and dose rates, but also the species and energy distributions of the particles composing the ionizing radiation. In LEO missions these data have already been extensively measured aboard robotic spacecrafts or the International Space Station. In the case of BLEO missions, a satisfactory amount of relevant data is also available as a legacy of many robotic missions flown in the inner and outer Solar System. A review of doses, particle content and energy distributions of SEPs measured in space is contained in [1] and references therein. In the case of measured Galactic Cosmic Radiation (GCR), [7] and [8] contain useful accounts and references. The analysis of data recorded by the RAD detector aboard the *Mars Science Laboratory* (MSL) while on route to Mars, showed [9] that the inclusion in the calculations of ambient radiation due to pions and their decay products makes an important difference when estimating the dose present behind spacecraft shielding. Information on relevant pion transport codes is available in [10, 11]. A quite detailed description of the radiation environment on Mars can be found in [12].

The second step of the present research consists in evaluating the risks related to the so-called deterministic effects. To this aim we use the formulas derived in [13], where sophisticated statistical methods have been applied to the analysis of events like the bombing of the Japanese cities of Nagasaki and Hiroshima or accidents involving radioactive substances. It is important to keep in mind that the parameters characterizing the exposure to radiation of the human population in such events are quite different from those describing the exposure in BLEO missions. For that reason, the coefficients computed for various types of deterministic effects in [13] are not entirely suitable in the case of space explorations. Nonetheless, it turns out that during a BLEO mission the risks of the appearance of any of the deterministic effects considered in [13] are so low, that they can be neglected even taking into account a large indetermination in the coefficients. This is not enough however to rule out deterministic effects. On the contrary, they may become relevant in the case of large SEP events, in which case significant acute doses are delivered to astronauts.

The risks related to the insurgence of stochastic effects like cancer are much more difficult to be estimated. Basing ourselves on the excess relative risk (ERR) for the induction of fatal cancer computed from the data of the Hiroshima and Nagasaki bombings, we estrapolate the ERR for a typical mission on Mars. These are rough estimations. More precise and refined calculations of the risks related to ionizing radiation are available like the NASA Space Cancer Risk model previously mentioned.

Despite many efforts, see [14] for a brief report on some of the conclusions coming from animal models and in vitro measurements, it is difficult to determine safe career dose limits for astronauts exposed to BLEO conditions and new strategies should be found. We propose an approach based on the so-called 3D in vitro models. These models, in which human organs are simulated by chimera organoids, have been widely applied in medicine in order to investigate new methods for curing cancer. They have also been used to track the initial steps leading to tumorigenesis in individuals affected by particular mutations. We propose to adopt a similar strategy to study the onset of cancer following the exposure to ionizing radiation, especially in the case of radiation containing heavy nuclei accelerated at high energies. This kind of radiation is in fact one of the major challenges to the human exploration of space, see for instance [15]. The idea of investigating the effects of radiation with the help of 3D models is not entirely new, see e. g. [14]. Strategies based on miniorgans (organoids) present several advantages, like for example the possibility of studying directly human cells avoiding the limitations of animal models in this respect. Moreover, organoids are now able to reproduce many of the characteristics of the original organs and even of the surrounding epithelium and supporting mesenchymal stroma. The challenge comes from the fact that, even weakening organoids by mutations or by infecting them with bacteria in such a way that they will be more likely to develop cancer, the probability of tumor induction following irradiation could be still too low to be observed. Besides, cancer is a multistep process that in humans can last for decades. We are however convinced that these difficulties may be overcome using the recent results of precision oncology and molecular medicine, in which the genetic and epigenetic changes leading to genomic instability and cancer formation have been pinpointed for several types of tumors. The revolution in understanding tumorigenesis in humans introduced by molecular medicine is illustrated here using the example of colorectal cancer, which has been extensively investigated in the recent past. We argue that some of the effects of radiation discussed in [14] have a deeper interpretation in the context of precision oncology.

### Three case studies: Missions to the Moon, Mars and asteroids

#### Doses and particle composition of the ionizing radiation in the case of missions to the Moon

We base our estimations on the extensive experience made during the Apollo missions, see [16]. The particle composition in that case included electrons, protons, neutrons, alpha particles and heavy ions. The radiation sources were provided by: particles trapped in the Van Allen belts (mainly energetic electrons and protons); Galactic Cosmic Rays (GCR); Solar Energetic Particles (SEPs) and secondary radiation created by the interaction of the primary particles with the spacecraft hull. According to [16], the doses received by the crews of the Apollo missions were significantly lower than the yearly average of 5 rem (50 mSv) set by the U.S. Atomic Energy Commission for workers exposed to radioactive sources. The main reason of these low levels is that during Apollo missions 7 – 15 no significant solar event associated with the emission of SEPs took place. The next two Apollo missions (16 –17) flew when the solar activity was at a minimum and the likelihood of large SEP events was accordingly low. We recall that Apollo 17 was the last human BLEO mission to be launched up to the year 2023. In all Apollo missions 7 – 17, the main source of ionizing radiation was GCR, the radiation levels of which are at their maximum during solar minimum. The estimated dose absorbed by each crewman on Apollo missions 7 – 15 averaged over the period of the travel from the Earth to the Moon and during the orbit around the Moon was of the order of 10 μGy/hour. In the period of ten days typical of Apollo missions to the Moon the resulting absorbed dose was of the order of 2.4 mGy [16]. On the lunar surface this value was less, i.e. ∼ 6 μGy/hour [16]. In this case, part of the absorbed radiation was due to neutrons produced in the collision of primary GCR with the lunar soil. The absorbed doses in Apollo missions 16 – 17 were higher, i.e. about 7.7 mGy [17], because these missions occurred at a time of solar minimum. More recent measurements made by the Cosmic Ray Telescope for the Effects of Radiation (CRaTER) instrument on the Lunar Reconnaissance Orbiter in the absence of SEP events, recorded a total absorbed dose of about 3.8 mGy in a ten days period. It is noted that the shielding of CRaTER (32 mils of aluminum, 1 mil=0.0254 mm) is comparable with that installed on the Apollo spacecraft (39 mils). The reasons of the discrepancy in the data of the absorbed dose measured during solar minimum between CRaTER and the Apollo missions 16 – 17 are explained in [17].

Summarizing, the absorbed doses that will be taken into consideration in this study with regard to missions to the Moon are the following:

1. an absorbed dose of about 10 mGy in the absence of SEP events
2. an absorbed dose of 4 Gy, which corresponds to the maximum operational dose (MOD) limit for skin that was set for the Apollo missions. This dose is comparable with that of 3.6 Gy that were estimated to be absorbed by the skin in the case of a large solar event [16].
3. an absorbed dose of 0.5 Gy for the blood forming organs set as a MOD limit in the Apollo missions, which is comparable to the estimated dose of 0.35 Gy incurred during a large solar event [16].

### Doses and particle composition of the ionizing radiation in the case of missions to Mars

The doses to which a human crew will be exposed during a journey to Mars and back can be roughly estimated thanks to the many unmanned missions to that planet. According to the data of the RAD detector on board of the Curiosity Rover, during the cruise phase to Mars of the Mars Science Laboratory mission, an average absorbed dose rate of:

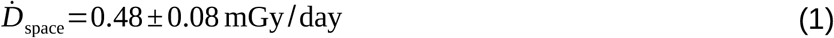

composed of GCR (mainly protons and alpha particles), was observed [18]. The average quality factor of GCR radiation measured in space was:

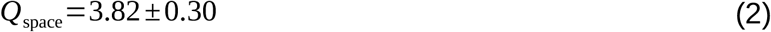

On the surface of Mars, the average absorbed dose rate amounted to

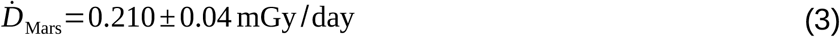

The average quality factor of the radiation on the surface (primary GCR plus neutrons and gamma rays coming from the secondary radiation) was:

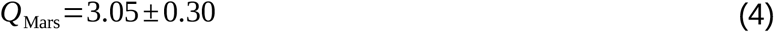

The SEP events registered during the Mars Science Laboratory mission consisted predominantly of protons for which *Q∼* 1.00 [18]. The delivered absorbed dose ranged from 1.2 mGy/event up to 19.5 mGy/event. In total, five SEP events were recorded during the Cruise Phase with an absorbed dose rate peak of more than 2.5 mGy/day. The total equivalent dose estimated in [19] for a cruise to Mars and back of duration 180 *×* 2=360days was 662 *±* 108 mSv. The contribution due to a stay on the Martian surface of 500 days was 320 *±* 50 mSv.

In total, this gives an equivalent dose of:

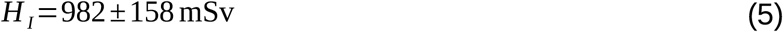

The corresponding absorbed dose, computed taking into account the quality factors *Q*_space_ and *Q*_Mars_ of Eqs. (2) and (4) respectively, is given by *D*_*I*_ =227.5 *±* 34.40 mGy. Considering instead a typical Mars mission with a short surface stay (consisting of 400 days in deep space and 30 days on the surface – as described in [1], the equivalent dose extrapolated from the above data is:

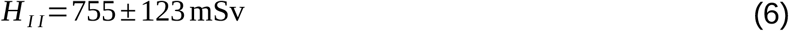

The related absorbed dose amounts to *D*_*I I*_ *∼* 198 *±* 33 mGy.

The total values of 982 *±* 158 mSv and of 755 *±* 123 mSv are both very high and comparable with the highest reference level recommended by ICRP for the rescue of workers on Earth in emergency situations.

### Biological response of humans to energetic particle irradiation Beyond Low Earth Orbit (BLEO) Deterministic effects

Deterministic effects are medically diagnosable effects related to cell killing caused by the absorption of ionizing radiation. These effects appear when the absorbed dose is high – of the order of several Gy ‘s – and affect single organs or tissues. The severity of the damage is dose dependent. In order to take into account the diversity in the response of individuals to ionizing radiation, it is possible to define lower and upper absorbed dose thresholds for the onset of a specific deterministic effect. The lower threshold *D*_*low*_ is such that the deterministic effect certainly does not appear for doses *D* such that *D* < *D*_*low*_. The upper threshold represents the dose at which the cell killing is high enough that the effect becomes unavoidable when *D* > *D*_*up*_. These thresholds depend on the way in which the dose is delivered. Their value is lower in the case of a single high dose delivered in a short time and higher when the dose is split into many smaller doses delivered over a long period. The risk *R* that a deterministic effect will appear in a sufficiently large sample consisting of individuals who have received a certain dose *D*, can be modeled using a sigmoid function. If *R* is defined as the percentage of individuals in which the effect becomes manifest, then the sigmoid function should approach zero when *D∼ D*_*low*_ and be nearly equal to 1 if *D∼ D*_*up*_ . For practical purposes *D*_*low*_ can be defined as the absorbed dose for which the risk of a given effect is *R* ( *D*_*low*_ )=0.01 [13]. Analogously, *D*_*up*_ is defined as the dose for which *R* ( *D*_*u*_ )=0.99. A possible choice expressing the dependence of *R* on *D* is provided using a Weibull function, which is of the form [13]:

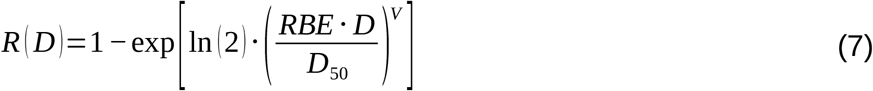

The number *N(D)* of affected individuals is given by the formula *N* ( *D* )=*N* _0_ *⋅ R* ( *D* ) where *N*_*0*_ is the total number of irradiated individuals.

The shape of the distribution in Eq. (7) is determined by three parameters:

- The absorbed dose *D*_50_ at which 50% of the exposed individuals are supposed to develop the given deterministic effect under study. This parameter depends on the dose rate according to the equation:

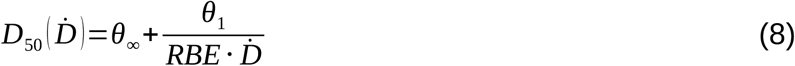

The first term in the right hand side of the above equation takes into account the case of acute dose rates delivered in a few minutes, while the second term, which is proportional to *θ*_1_, is the correction required by the fact that the effects of radiation vary as a function of the dose rate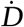. It is pointed out that Eq. (8) is purely phenomenological. The parameters *θ*_*∞*_ and *θ*_1_ are extrapolated applying sophisticated statistical methods to the existing data from accidents involving radioactive sources.

- The relative biological effectiveness factor *RBE*.
- A shape factor *V* determining the slope of the sigmoid. By adjusting *V*, it is possible to accommodate different values of the lower and upper thresholds *D*_*low*_ and *D*_*up*_. Examples of risk-dose curves are given for various values of the parameter *V* in Figure 1. As in the case of *θ*_*∞*_and *θ*_1_, *V* is evaluated on the basis of statistical methods.

**Figure 1.**
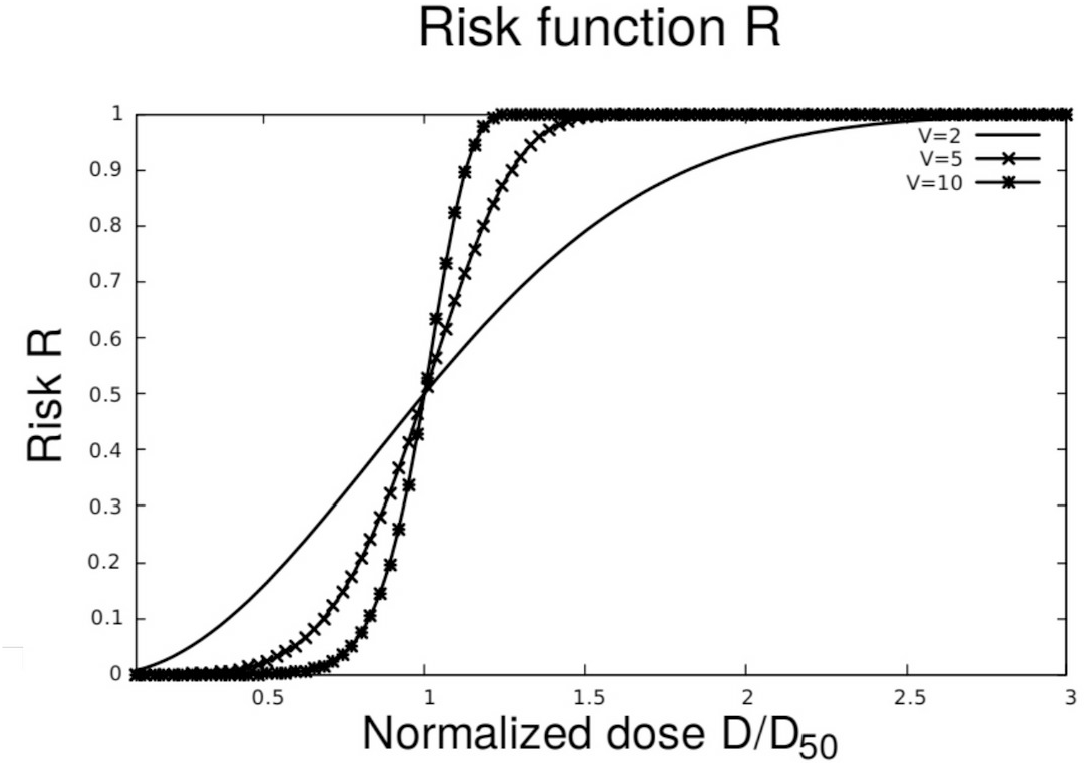
Risk – dose curves computed using the sigmoid function defined in Eq. (7) for various values of the parameter *V*.

The parameters *θ*_*∞*_, *θ*_1_and *V* are effect dependent.

### Deterministic effects in the case of Mars missions

We are now in the position to study the risk of the onset of deterministic effects during a space mission. First we discuss the case of Mars missions. Equation (7) only provides indicative results. To make more quantitative predictions, it would be necessary to have a sufficient statistics regarding the action of ionizing radiation of space origin in producing deterministic and stochastic effects in the human body. Clearly, this statistics is not available in the case of flights to Mars. All existing formulae that attempt to extrapolate the risk of mortal or morbidity effects are based on data coming from nuclear accidents on the Earth. These formulas are forcefully affected by several uncertainties – for instance in determining parameters like the RBE and the factor *D*_50_ defined in Eq. (8) or, in the case of stochastic effects, the dose and dose rate effectiveness factor (DDREF). The uncertainties in the DDREF are supposed to become particularly large when the dose rates are lower than 0.05 Gy/h [20]. The dose rates expressed in Eqs. (1) and (3) which are relevant to Mars missions, are much lower than that value, even taking into account the occurrence of a large SEP.

The coefficient *D*_50_ may be computed from Eq. (8) using the parameters *θ*_1_, *θ*_*∞*_ and *V*. The values of the RBE needed for the calculations are approximated starting from the radiation quality factor for space *Q*_space_ given by Eq. (2). The use of the quality factor, which refers to the induction of stocastic effects, is a crude approximation made necessary by the fact that the RBE related to a given deterministic effect due to radiation in space is not known. It is however a reasonable approximation: The International Commission for Radiobiological protection recommends the use of the weighting factors for stochastic effects as a conservative criterion to predict the risks of deterministic effects except in special cases in which high-LET radiation is the critical factor [21]. Assuming further that the stay on the surface of Mars is short, of the order of 30 days as it is typical of a short-term mission [22], we choose as effective dose G for the whole mission *G*=*RBE ⋅ D* a value that is about four times larger than the values *H*_*I*_ and *H*_*II*_ coming from the data of the Curiosity mission given in Eqs. (5) and (6) respectively:

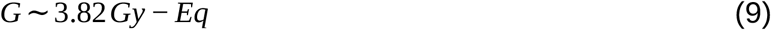

This value, given in Gray-Equivalent units, is big enough to include an almost 400% uncertainty related to the rough approximation used in deriving the coefficient RBE. It also allows to consider a worst case scenario characterized by a large SEP event. The increase of the dose of 400% is justified by the comparison of the quality factor and the RBE for deterministic effects in the case of some HZE particles, see [23]. Another quantity that will be relevant in the calculation performed in this subsection is the dose rate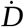 . A reasonable choice of 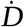 is the average absorbed dose rate in space given in Eq. (1). We rewrite it here using a slightly different set of units, that fits with the units in which the parameters *θ*_*∞*_ and *θ*_1_ will be defined afterwards:

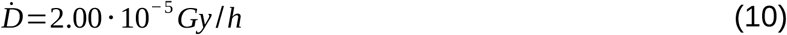

We suppose that there is no attenuation of the radiation delivered to an organ due to its absorption by the surrounding tissues and by the hull of the space vessel. This hypothesis can be justified as follows. First of all, we expect that the overwhelming contribution to the absorbed dose comes from highly energetic protons and we consider a stay on the surface of Mars of the order of one terrestrial month. In this case, the total absorbed dose will be delivered mainly in space and will predominantly consist of GCR. The main contribution to the particle content of GCR is made by protons (about 80%) whose energy spectrum peaks in the energy range between 300 MeV and 1 GeV [24]. Already at 300 MeV, the range of protons in aluminum is about 6.57 g/cm^2^, i.e. 24.34 cm, while at 1 GeV the range grows to 4.10 *⋅* 10^2^ *g* / *cm*^2^, corresponding to 152.60 cm. In consequence, the percentage of particles stopped by the hull of the spacecraft, even assuming a thickness of 24 cm of aluminium, is limited. In this situation, in which there is a predominance of light, very energetic nuclei, the doses related to secondary particles, in particular neutrons, can be considered to be negligible, at least to a first approximation. A fortiori, it is possible to neglect the difference among the doses absorbed by different tissues in the human body. As a matter of fact, the average density of human tissues is lower than that of aluminium (by about a factor 2.70) and their stopping power is proportionally less. For example, approximating the density of a tissue with that of water, it turns out that the range of protons at 300 MeV in water is about 5.14 g/cm2(*∼* 51.40 *cm*). At 1 GeV the range increases to 325 g/cm2 (*∼* 325 *cm*). Clearly, in this situation there is no attenuation of the radiation delivered to an organ due to its absorption by surrounding tissues. Apart from protons, another component of the cosmic rays that may have relevant health effects are highly energetic heavy nuclei. The energy spectrum per nucleon of these particles is peaked in the energy range between 300 MeV and 1 GeV [24]. These particles have a high penetrating power inside the human body. Though their flux is many orders less than that of protons, heavy nuclei can have disastrous consequences for the health of the crew of a manned mission ([20], [25]), mainly because, when hitting the nucleus of a cell, they produce localized multiple damage in the DNA, which are much more difficult to repair than the damage caused by low-LET radiation. The effects of heavy nuclei cannot be taken into account by Eq. (7) for reasons that will be explained later. Also, some of these effects, for instance on the functionality of the brain and of the central nervous system, will be discussed separately.

On the basis of Eq. (7) it is possible to exclude the occurrence of several fatal deterministic effects, like the gastrointestinal syndrome and the pulmonary syndrome discussed in [13]. These pathologies are simply beyond the lower threshold of occurrence *D*_*low*_. What cannot be entirely neglected is the risk *R*_*hs*_ of the hematopoietic syndrome (also called bone marrow syndrome), which amounts to:

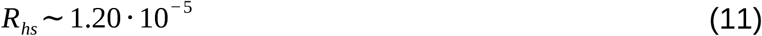

In this syndrome the killing of blood cell precursors in the bone marrow is so high that blood cells in the body cannot be substituted for them at the rate required. The main consequences are bleeding and a lower ability to fight infections for a long period after irradiation, see [13] and references therein for more details.

In computing *R*_*hs*_, the parameters *θ*_*∞*_, *θ*_1_, *V* are set following [13]: *θ*_*∞*_=3.00 *Gy, θ*_1_=0.07 *G y*^2^/ *h* and V=2. We recall that in the present subsection, the equivalent dose and the dose rate are given by Eqs. (9) and (10) respectively.

Morbidity effects seem not to be very likely. For instance, the risk *R*_*cat*_ of cataracts of the eye (*θ*_*∞*_=3.00 *Gy, θ*_1_=0.01 *Gy*^2^/ *h, V* =5)amounts to:

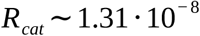

The risk of vomiting *R*_*v*_ is instead given by (*θ*_*∞*_=2.00 *Gy, θ*_1_=0.20 *Gy*^2^/ *h, V* =3):

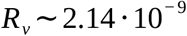

Relevantly higher is the risk of diarrhea (*θ*_*∞*_=3.00 *Gy, θ*_1_=0.20 *Gy*^2^/ *h, V* =2.50):

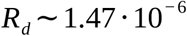

These differences are however not really significant. What is significant is that the order of magnitude of the values of all the risks of deterministic effects computed above are very low, implying that the occurrence of these effects is unlikely.

It is worth to stress that in the case of the central nervous system (CNS) syndrome, it is not possible to specify the parameters *D*_*50*_ and *V* with the statistical techniques used in [13]. The reason is that to determine the symptoms of this syndrome is not an easy task. Even when very mild symptoms are presented, a patient affected by the CNS syndrome will die within 48 hours due to several possible causes. According to [13], the CNS syndrome occurs at acute doses above 50 Gy. In the case of Mars missions, we have seen that the doses delivered are far below this threshold while also the dose rates are much lower than those typical of acute doses. Despite that, more subtle influences can be exerted on the CNS due to GCR, which do not lead directly to death, but constitute a serious danger of impairment for the crew members of a human BLEO mission. The reason is the presence in GCR of heavy nuclei accelerated to very high energies, the so-called HZE particles. Such particles are not efficiently stopped by the shielding of the space vessel and, when hitting a non-dividing cell such as a neuron, are very likely to compromise their functions in a relevant way. This problem has been known for many years. Already in 1973, an advisory panel to the Space Science Board of the US National Academy of Science urged a careful investigation of this aspect of GCR [26]. In [27] it was shown using numerical simulations that, during a three-year mission to Mars at solar minimum, HZE-particles may damage extended portions of a relevant percentage of hippocampal and thalamus cell nuclei. The effects of this radiation can be asymptomatic for a long time due to the exceptional abilities of the CNS to overcome damage. However, the authors of [27] did not exclude the related possibility of complete inactivation of the neurons and/or the onset of a series of events leading to cell killing or functional impairment. This subject has attracted considerable attention in recent times due to new results coming from experiments on mice. These results show that even relatively small doses of heavy nuclei (of the order of a few tens of cGy) lead to alterations in neurons and in synaptic connectivity, eventually causing a significant behavioral impairment in the irradiated mice ([28], [29]).

In conclusion, the probability of deterministic effects studied here with the help of the statistical methods of [13] is very low when doses typical of a mission to Mars are considered. The obtained results are in agreement with previous publications on the subject. For example, even in the case of acute doses of ionizing radiation, the threshold for the hematopoietic syndrome is > 1 Gy and generally there are no clinically significant effects if the dose is below 2 Gy, [30] and [31]. However, this does not mean that there are not other deterministic effects that could become relevant during space travel. An example are the damages to the CNS induced by HZE radiation. Moreover, it is known that during LEO space flights the crew is affected by disturbances related to the impairment of the immune system, see for instance [32], [33]. Nevertheless, due to the limited statistics available thus far, it remains to be proved if such disturbances have long-term consequences on the health of astronauts or on their lifespan^1^. In general, it is not easy to separate the short-term deterministic effects of ionizing radiation from those due to all the other factors that can impact the health state of a human crew in space. Examples of such factors are microgravity, isolation and emotional stress.

### Deterministic effects in the case of Moon missions

In the case of short missions to the Moon, like the Apollo sequence, it is possible to rule out deterministic effects, at least in the absence of a large SEP event. In fact, absorbed doses of the order of 10 mGy are far below the lower threshold *D*_*low*_ of any lethal or non-lethal deterministic effect. However, a prolonged stay in the van Allen belts could be dangerous. Assuming an equivalent dose rate of about 13.33 mSv/h [23], it is possible to use Eq. (7) in order to evaluate the lower and upper threshold doses for the onset of a given deterministic effect. For example, in the case of the bone marrow syndrome described above, the lower threshold dose is *D*_*low*_∼3.30 Gy, corresponding to a stay in the Van Allen belts of about 248 h. The upper dose *D*_*up*_ amounts to 21.50 Gy and the relative exposure time is about 1610 hours.

An important phenomenon is that of the light flashes which were perceived by individuals subjected to ionizing radiation mainly in dark environment conditions. This effect could be potentially dangerous if tasks should be performed which require a high level of visual efficiency. Further, light flashes could cause disturbances in the sleep cycle. According to data collected during the ALTEA mission [35], about 30 events were observed in the average by astronauts in the USLab module of the International Space Station (ISS). The rate of flashes varied depending on the shielding of the spacecraft and on individual physiological and psychological characteristics [35]. The identity of particles causing light flashes is still a subject of debate. In space, candidates are light ions with charges ranging from 3 *≤ Z ≈* 8, although heavier ions are also suspects [35]. The latter, like for instance *Fe*, are more difficult to investigate because they are less abundant in LEO than are lighter ions. On the ground also light particles like pions, muons and alpha-particles have been shown to cause light flashes [35]. More recently, cases involving high energy photons (6 MeV) have been reported [36]. The possibility that protons too could be associated with the phenomenon is under discussion [35]. The mechanism by which individuals whose eyes and/or cortex are irradiated perceive light in the complete darkness is not yet fully understood. According to [35], it is for instance not clear if there exist some mechanism to “see” ionizing radiation directly or if the observed light flashes are due to photons in the visible spectrum emitted after the high energetic particles hit an area near the retina.

## Stochastic effects

Stochastic effects can be somatic – cancer induction (leukemia or solid tumors) – or heritable effects (also called hereditary effects). In contrast to deterministic effects, it is believed that there is no threshold dose for the onset of the stochastic effects of ionizing radiation. More precisely, the probability of the onset of a stochastic effect, but not its severity, may be regarded as a function of the delivered dose [37]. According to this definition, every dose, no matter how small, is able to cause severe effects like cancer induction or mutations in individuals hit by ionizing radiation and in their next generations. Hereditary effects have proved to be very difficult to study in humans and, up to now, they have been predominantly tested on animals. According to [38], there is an emerging consensus that transmissible chromosomal instabilities do not occur in response to low-LET radiation below 0.5 Gy, implying the existence of a threshold at least in this case. This conclusion does not however apply to the high-LET radiation, relevant to space missions.

### Dose-response curves and low-high doses

Cells exposed to ionizing particles show a response to the dose of received radiation that is measurable. Possible effects can range from the molecular level, e.g. single and double strand breaks up to more complex cytological and genetic lesions, such as the formation of micronuclei, chromosomal aberrations and mutations – including changes that lead to cancer. At fixed doses, the response of living cells depends on many factors, the most important of which is probably the dose rate. In mice, for instance, an acute dose of the order of 10 Gy delivered in a short term causes severe illness and is mostly fatal, whereas the same dose delivered over a 450-day period is apparently harmless [39]. However, it should be taken into account that high accumulated doses may result in the occurrence of tumors after many years, an outcome which makes the study of stochastic effects very difficult.

Other relevant factors that determine the response of cells or tissues include the type of radiation, as well as physiological and environmental conditions. The concepts of low dose and low dose rates have been extensively discussed in the literature, see e. g. [40] – Annex G, using epidemiological studies on populations affected by radiation emitted during the Japan bombings or nuclear accidents and taking into account as an effect solid tumor induction in humans. A low dose has been defined in [41] to be any exposure below *D*_*max − low − dose*_ *∼* 200 mGy^2^.

At the moment, there is no satisfactory model which is able to predict the possible mutagenic, cancerogenic and teratogenic effects of radiation at low doses. The standard linear no-threshold model (LNT model), which assumes that the risk of such effects per unit dose is independent of the dose delivered, is currently a subject of debate. Several alternatives to correct this model at low doses are discussed in [43]. The results of many experiments aiming to determine cell damage at acute low doses can be found in [44]. See also [45] for a review of epidemiological studies. The investigation of low doses is not entirely relevant in the case of space missions, because the doses involved are much higher. Yet, some interesting related phenomena like the Bystander Effect [46] and the Adaptive Response may nevertheless be of some interest. The peculiar situation of space missions, in which intermediate and high doses are delivered at relatively low dose rates, certainly deserves more attention. Studies of this kind, especially in the case of high-LET radiation deposited by heavy ions, are still scarce.

### An approximate calculation from statistical data assuming the linear no-threshold model

Assuming that the dose-response curve for radiation-induced cancer follows a linear model with no threshold, we will provide an estimation of the probability of the onset of stochastic effects under space conditions using the statistical data coming from low-LET radiation. These predictions should be considered as qualitative. For example, they do not take into account several factors that are able to change the response of living organism to ionizing radiation, such as for instance the bystander effect, and the action of repair mechanisms in aerobic/hypoxic conditions. It should also be stressed that nowadays very sophisticated models can be used in order to predict the risks of ionizing radiation. Examples of these models are for instance the BEIR VII risk models worked out by the BEIR VII Committee [47]. While their detailed discussion is out of the scope of this work, the risk models in [48] will be briefly mentioned below.

One of the main ingredients of our considerations is the excess relative risk per unit dose *ERR*_*Z*_. This quantity measures the percentage increase of risk of contracting fatal cancer in a group of *N*_*0*_ individuals exposed to radiation of type *Z* with respect to a group of *N*_*0*_ unexposed individuals. In the case of the atomic bombings in Japan, the statistical analysis of the data obtained from the survivors has shown that the excess relative risk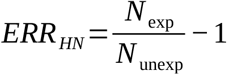 for the induction of fatal cancer per unit dose is^3^

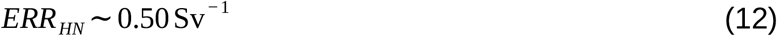

where the subscript *HN* refers to the Japanese cities of Hiroshima and Nagasaki. Moreover, *N*_exp_ is the number of events occurring in the group of *N*_*0*_ exposed individuals, while *N*_*unexp*_ denotes the number of events in the control group consisting of *N*_*0*_ unexposed individuals. ‘An event’ means here a mutation leading to fatal cancer. A crude estimate of the excess relative risk per unit of dose in space *ERR*_*space*_ during the journey to Mars can be obtained using the fact that the excess relative risk scales roughly according to the quality factor *Q* of the radiation type. We notice that a similar strategy based on the dose equivalent H=QD has been recently endorsed by the National Academies of Sciences [50]. Thus, denoting with the symbol *Q*_*HN*_ the quality factor of the radiation in the bombing of Japan, an approximate evaluation of *ERR*_*space*_ is:

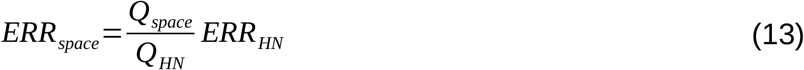

The quality factor *Q*_*space*_ is already known from Eq. (2). To determine *Q*_*HN*_, we proceed as follows. The radiation due to the atomic bombs at Hiroshima and Nakasaki consisted not only of photons, which are low-LET radiation, but also of neutrons with a quality factor of about 10. Following [49], we assume that the neutrons constituted 1% of the total radiation^4^. In consequence, the quality factor *Q*_*HN*_ is approximately:

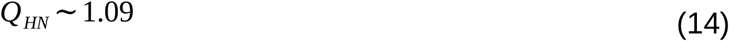

Substituting this value in Eq. (13), we obtain:

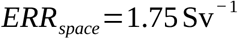

This value is quite high. It implies that the probability of fatal cancer during the entire life (about 20% in the unexposed population) raises to 55% after an exposure to a sievert of space radiation during the travel to Mars.

Based on the *ERR*_*Z*_ and other similar quantities, it is possible to establish models that attempt to assess the risks connected to irradiation. For instance, let *ERR*_*Z*_*(D,e,s)* be the excess relative risk of contracting fatal cancer after an exposure at the age *e* of an individual of sex *s* to a dose *D* of radiation of type *Z*. The so-called probability of causation, in short *PC*, estimating the chances that the absorbed dose of radiation has been the cause of fatal cancer after the exposure, may be approximated as follows [48]:

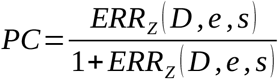

The formulas for evaluating *ERR*_*Z*_*(D,e,s)* may be found in [48]. Another relevant quantity is the Risk of Exposure Induced Death (REID). The REID accounts also for cancer deaths that would occur anyway, but are shifted to an earlier age due to radiation exposure. The NASA career dose limits in sieverts are established by requiring that the exposure to radiation during the career of a particular astronaut is not to exceed 3% REID from fatal cancers at the upper 95% confidence level, see for instance [51]. More information on the evaluation of the risks of stochastic effects due to ionizing radiation can be found in [2]. A new approach to reduce the uncertainties on parameters like the quality factor of radiation in space has been presented in [52].

### Need of a new strategy to ascertain the safety of BLEO missions Statement of the problem and outline of its possible solution

Due to the lack of sufficient statistics it is difficult to predict the insurgence of deterministic, stochastic and heritable effects. An additional complication for BLEO missions is provided by the fact that the radiation contains a relevant component of HZE particles whose effects on humans are still largely not understood [20], [53-55].

A comprehensive review on the state of our understanding of the effects of HZE particles exposure can be found in [14]. In that publication, the phenomenon of tumorigenesis after HZE particle exposure is tackled by considering the relation between the risks of cancer induction and the oxidative stress following irradiation. An extensive series of interesting experiments are reported which show how the production of reactive oxygen species (ROS) after a dose of HZE radiation may last for years, interfering with the cell repair system, affecting telomere stability and favoring chronic inflammation. It is also hinted that ROS might affect the redox regulation of proteins and lipids in cells, changing in this way the epigenetic landscape, a fact that could lead to heritable effects. The recommendation of [14] is that, in order to understand the phenomenon of tumorigenesis, it is critical to evaluate how the persistent production of ROS and their moieties depend on the charge, energy and LET of the imparted ionizing radiation. The work of [14] presents also the state-of-the-art of in vivo experiments performed on mice models in order to assess the risks of cancer related to ionizing radiation in humans.

One of the conclusions coming from these studies on mice is that the risks after irradiation of the insurgence of a number of different types of carcinomas is largely dependent on LET. It is also possible that there exist LET thresholds for the induction of certain kinds of tumors. This seems to be for instance the case for the thymic lymphoma investigated by [56].

Despite all advances, which significantly improve our understanding of the risks of HZE radiation, several problems in determining safe career dose limits for astronauts still persist. Limitations in the prediction of the health risks incurred due to ionizing radiation in space are discussed in [15] and [57]. The key point is that with experiments in vivo or in vitro^5^ it is extremely difficult to determine the probability of stochastic events that, in humans, may require decades before they occur. To this purpose it is of uttermost importance to determine and apply markers that are able to signal in advance the early stages of carcinogenesis in humans. The human genomic features associated with a higher likelihood of developing given types of cancer should also be taken into account.

A new strategy to overcome at least in part these difficulties and to achieve the goal of establishing career limit for astronauts engaged in BLEO missions with a safety level comparable to that NASA criteria for LEO missions [51] could come from some recent results of precision oncology and molecular medicine. In particular, thanks to an extensive effort in clinical testing supported in the laboratory also by the application of human organoids cultures and genome editing technology, it has been possible to characterize with a high precision the pathways of tumor formation in selected human organs. This knowledge, combined with the use of organoids could be helpful in solving the problems mentioned before by integrating the data coming from in vitro experiments and animal models. First of all, the possibility of detecting the early stages of cancer in an organ only by checking the activation of certain oncogenic pathways (such as the TGF-β signaling pathway), the loss of heterozygosity or other gene alterations, allows to shorten the time for predicting if there are risks of cancer induction due to irradiation. If required, it is even possible to transplant tissues from these organoids at any stage of the evolution of their health after irradiation and to check their level of tumorigenicity. Secondly, organoids can be formed using human cells, thus avoiding the limitations due to the fact that in animal models the cells can have a different response to radiation. The growth of organoids is technically more challenging than that of 2D models of cultured cells. Yet organoids still retain the ease of maintainance and analysis typical of in vitro models and, in addition provide in the context of the study of the early steps of the onset of cancer a much more refined simulation of human organs than traditional cultured cells. It is sufficient here to mention the fact that cells in a 2D culture lose their differentiation in most cases ([58], [59]) and their polarity. Both features are preserved when cells are put in a 3D environment [60], and sometimes can be restored when passing from 2D to 3D [61], [62]. Gene expression and reproduction are also very different than in vivo [63]. For instance, the loss of polarity affects also the intracellular signalling pathways [63] that, as we will see later, play a crucial role in the development of cancer.

The present state-of-the-art of our knowledge of the early stages of cancer thanks to precision oncology will be illustrated in the next Section taking as an example the specific case of colorectal cancer. Next, the applications to space medicine outlined above will be explained in more detail.

### Colorectal cancer as a study case

The events leading to colorectal cancer (CRC) in humans have been satisfactorily understood at the molecular level. This makes it possible to detect the early stages of colorectal carcinogenesis with the help of techniques that are already commonly applied like RNA-sequencing and Loss of Heterozygosity (LOH). Comparable knowledge already exists or is soon to be obtained for the case of solid tumors affecting other organs, like for instance pancreatic cancer [64]. All of these advances together with the use of organoids open the possibility to achieve a better understanding of the risks of stochastic effects related to exposure to ionizing radiation without the need of waiting decades for the appearance of such effects in vivo or reliance on complex epidemiological studies.

Coming back to colorectal cancer, its development follows a pathway of ordered events transforming normal epithelium cells into adenoma cells and finally leading to adenocarcinomas. “Driver” events are certain mutations – later discussed in details - activating oncogenes, tumor suppressor genes and genes responsible for activating certain signal transduction pathways that regulate the response of cells to stresses. Molecular studies of precision oncology have revealed that these driver events are not sufficient to initiate tumor progression. The presence of genomic instability is required in order to increase the rate of mutation necessary for cancer formation. Remarkably, the set of driver events and genomic instabilities that are relevant for colorectal cancer is limited, a fact that potentially allows verification in the laboratory whether, at some stage after irradiation, the conditions for the formation of a tumor are present or not. Out of 80 genes that are involved, it has been found that the real drivers consist of mutations of a much smaller number of genes [65]. The genomic instabilities involved are only three. With regard to driver events, relevant is the inactivation of the APC (Adenomatous Polyposis Coli) tumor suppressor gene followed in the majority of CRC ‘s by the mutation of the KRAS gene, a proto-oncogene that regulates cell growth. More rare is the mutation of the BRAF proto-oncogene.

It is noted that the mutations of the KRAS and BRAF genes are mutually exclusive. The coexistence of both mutations has been observed in only 0.001% cases of CRC. Other important genes in KRAS mutated CRC ‘s are the TP53 and SMAD4 tumor suppressor genes and the PIK3CA oncogene^6^. In BRAF mutated CRC ‘s affected are genes related to the DNA mismatch repair system like MLH1, MLH3, MSH2, MSH3, MSH6 and PMS2. These genes are usually not mutated, but rather epigenetically silenced, although their mutation is possible, but occurs less frequently. The signaling transduction pathways that are altered in CRC are those that regulate cell differentiation, migration, proliferation and apoptosis. In particular, the WNT-β-catenin and the transforming growth factor-β (TGF-β) pathways are affected. For instance, the APC gene synthesizes a large protein with multiple functions including apoptosis and cell differentiation and interacts with the β-catenin. The latter is a protein that regulates the adhesion of cells as well as gene transcription. It is noted that the activation of the WNT pathway is connected to an increased production of ROS that cause further molecular damage [66]. The WNT pathway is activated in 93% of CRC ‘s [67]. The role of ROS is one of the key points for understanding cancer development after exposure to radiation, see [14]. As we see, the results of molecular medicine provide a more fundamental insight into the mechanisms that cause oxidative stresses in cells. The other crucial affected pathways are TP53, PI3K, the transforming growth factor beta (TGF-β) and MAPK [67], Epithelial-to-mesenchymal transition (EMT) and MYC oncogene amplification. The gene mutations in the early development of CRC take place at a fast rate that cannot be explained by the random mutation rate occurring in normal cells. As already anticipated, for initiating cancer the presence of genomic instabilities favoring the accumulation of mutations is necessary. While it is not clear if the genomic instabilities start the mutations or vice-versa, the three types of genomic instabilities leading to CRC are known. In about 85% of CRC ‘s, the driving events are accompanied by chromosomal instability (CIN), while the remaining 15% are characterized by microsatellite instability (MSI) [68]. In tumors with MSI, the occurrence of a high CpG island methylation phenotype (CIMP) is frequent [68]. Basically, CIN can be related to double strand breaks with subsequent defective repair leading to a fast rate of gain or loss of large portions of chromosomes, a phenomenon that causes somatic copy number alterations (SCNA) of genes. More precisely, CIN arises from defects of chromosomal segregation and of telomere stability [68]. The CIN condition is also favored by mutations in the TP53 pathway and other crucial genes [68]. The alterations related to CIN change the number and appearance of the chromosomes, in short, the karyotype of cells. While normal human cells are diploid, i. e. chromosomes are present in two copies, cells affected by CIN can have chromosomes with a different number of copies and some of their chromosomes may suffer the loss of heterozygosity (LOH). LOH occurs when one gene in one allele is lost. In cancer this becomes a crucial event when the gene is a tumor suppressor and the copy of that gene in the other allele is inactive due to a point mutation. The gain or loss of large portions of chromosomes usually ends up with the shortening of telomeres^7^. The CIN-pathway to tumorigenesis can be summarized as follows. With the activation of the WNT signaling pathway, whose proteins decrease the stability and levels of the proteins devoted to the degradation of β-catenin, the latter protein starts to accumulate in the cell cytoplasm. The damage of tumor suppressor genes like the APC gene, promoting the degradation of the β-catenin protein, contributes to this accumulation. The β-catenin proteins are responsible for cell growth and cell adhesion. When they translocate to the nucleus, one of their actions comprises interfering with the transcription of the MYC oncogene. The mutation of the KRAS gene further promotes cell proliferation and inhibits cell apoptosis. The SMAD2 and SMAD4 tumor suppressor genes also mutate inhibiting the TGF-β pathway. This leads to uncontrolled cell proliferation. At a later stage the mutation of TP53 also occurs. As already said, this fast rate of mutations becomes possible due to CIN. The techniques used to detect CIN are described for instance in [65]. They include cytometry, karyotyping, loss of heterozygosity, fluorescent in situ hybridization (FISH), and comparative genomic hybridization (CGH). The second pathway to genomic instability, MSI, is related to gene loci (the microsatellites) characterized by sequences of one to six nucleotides, for instance (AC)_*n*_, that are repeated a number of times *n* ranging from five to one hundred. The human genome contains about 50000 – 100000 microsatellites that are found mostly in the non-coding regions of DNA, but also in the coding part. The presence of repeated units creates motifs in the DNA that are used as markers for various enzymes, although most microsatellites look to be neutral markers. As markers, microsatellites play a role in DNA recombination and replication. Moreover, they influence gene expression. More details concerning their functions can be found in [71]. Microsatellites are areas intensively subjected to mutations because their regularity is easily altered during recombination and replication. For this reason, they are very sensitive to errors in the DNA mismatch repair system Methyl Mismatch Repair (MMR), which is responsible for the recognition and repair of mistakes occurring when DNA replicates or recombines. The proteins involved in MMR are coded by seven genes: MLH1, MLH3, MSH2, MSH3, MSH6, PMS1 and PMS2. The mutation or epigenetic silencing of some of these genes leads to MSI. The MSI pathway to cancer can be summarized as follows. Microsatellites are found in the powerful tumor suppressor gene TGF-β that regulates cell division. The malfunctioning of MMR leads to MSI and favors the further stages of carcinogenesis, accumulating mutations in genes responsible for cell differentiation, apoptosis and growth. The alteration of the TGF-β pathway with the subsequent coding of abnormal typeII TGF-β receptor and BAX protein leads to cell proliferation^8^. The mutation of the BRAF gene is an important event in MSI tumorigenesis too. Let us also note that the mutations of TP53 and of the ATM gene are mutually exclusive. The alteration of ATM gene is predominant in the MSI phenotype. In MSI driven CRC a relevant role is played by epigenetic silencing of the critical genes. For instance, MLH1 is epigenetically silenced by CIMP (CpG Island Methylator Phenotype). CIMP refers to the methylation of the CpG islands. These are regions of DNA in which there is a high frequency of CpG sites. Such sites consist of a cytosine nucleotide connected to a guanine nucleotide by only one phosphate. Formally, a CpG island is defined as a region of at least 200 base pairs in length in which the percentage of dinucleotides CpG is greater than 50%. The CpG islands in humans and, more in general, in vertebrates, are located in the promoters or near the promoters. In humans, CpG islands are present in about 70% of promoters. Promoters can be found in the initial part of genes and play an important role in transcription. As a matter of fact, to start the gene transcription, the enzyme that synthetizes RNA, known as RNA polymerase, must bind to the promoter. If the binding site is inactivated, the gene is no longer recognized by the RNA polymerase and is not expressed. In CRC tumor suppressor genes and other genes related to tumor development are epigenetically silenced by hypermethylation of the CpG islands. Methylation is a process in which the cytosine of the CpG sites has a methyl group attached. In CIMP, the CpG islands in the promoters of multiple genes are simultaneously hypermethylated. For the analysis of MSI several methods may be applied, like immunohistochemical (IHC) staining and PCR amplification. The IHC analysis is performed using antibodies to identify the loss of proteins that are involved in MMR. In the PCR analysis a set of suitable microsatellite markers is used, like for instance the BAT25, BAT26, D2S123, D5S346 and D17S2720 markers. For instance, D2S123 is a microsatellite linked to the MHS2 gene containing repeat of CA (cytosine – adenine) with chromosomal location 2p16. Concerning CIMP, up to now there is no a consensus methodology to check CIMP status [73]. Different panels of markers used by independent groups can be found in the review [73].

To conclude, the progresses of molecular medicine are not only restricted to CRC. Similar results have been obtained also in the case of other cancers, like for instance hepatocellular cancer, biliary cancer and pancreatic cancer [74]. Of course, it should not be forgotten that the research in this subject is still ongoing. In particular, a unique classification of the subtypes of CRC with respect to their response to given therapies still does not exist. Depending on the markers and methodology used, different groups have provided several classifications of CRC in which the number of subtypes varies from three to six. We report here the four consensus molecular subtypes (CMS) that have been distinguished after an extensive analysis of 4000 samples of CRC [75], see also [68] for a review:

CMS1: This subtype (14% of early-stage tumors) is characterized by MSI and hypermethylation of the CpG islands (CIMP). The level of CIN is low. Often the mutation of the BRAF gene is observed. CMS1 is prevalent in lesions appearing in the right colon [67]. In CMS1 cells are hypermutated^9^.

CMS2: This group (37% of early-stage tumors) is microsatellite stable, while CIN is marked, but not so dominating as in CMS4. There is a strong upregulation of the WNT and MYC pathways^10^. Moreover, the expression of the oncogenes EGFR, ERBB2 is higher or amplification of these genes occurs. These events are accompanied by mutations of the gene TP53. CMS2 is preferably diagnosed in the left colon and rectum [67].

CMS3: This subtype (13% of early-stage tumors) belongs to the CIN group like CMS2 and CMS4 but has distinctive characteristics. In particular, the level of CIN is low and the upregulation of the WNT and MYC pathways is moderate. With respect to CMS2, KRAS mutations are enriched. There are mutations in the genes belonging to the PI3K pathway. In this kind of CRC the most important feature is the ability of cancer cells to undergo metabolic reprogramming. About 30% of these tumors have MSI, CIMP and are hypermutated. This subtype appears in the large intestine with no preferred location.

CMS4: The CMS4 subtype (23% of early-stage tumors) is characterized by TGF-β activation and by upregulation of genes implicated in the epithelial-mesenchymal transition (EMT). With respect to epithelial cells, mesenchymal cells have migratory and invasive properties that worsen the prognosis for this kind of tumor. A better characterization of CMS4, in particular the knowledge of the driver events leading to CMS4 and the regulatory mechanisms underlying the expressions of the genes involved, can be found in [77].

CMS2-4 follow the CIN phenotype described before in this Subsection, while CMS1 follows the MSI phenotype. CMS1 is typically hypermutated, while CMS2-4 is not, with exceptions in the case of CMS3. The mutation distributions of different genes within the consensus molecular subtypes and other molecular data that help in distinguishing these groups can be found in [78]. The consensus molecular subtypes have been obtained through combining the efforts of different research teams and they provide the most robust classification of CRC available thus far. Still, 13% of CRC do not fit in any of the four CMS groups presented above. Research is still ongoing in this regard and the attainment of a more complete understanding of the molecular event, markers and pathways is necessary before it will be possible to develop efficient drugs and therapies to cure CRC. Deficiencies in the present knowledge of CRC certainly pose a limitation for clinical purposes, but not for astrobiological purposes. In fact, the progress mentioned above provides powerful criteria to support recognition of the first stages of cancer after irradiation as well as concrete tests for checking the driving mutations of the critical genes, the presence of genomic instability and of epigenetic alterations. In the next Subsection, the application of these ideas to the case of organoid cells will be proposed.

### Organoid technologies helping in establishing safe career dose limits for BLEO missions

Organoids, also called 3D in vivo cell cultures^11^, are multicellular structures that are able to mimic some of the characteristics of in vivo organs. They can be obtained starting both from adult cells or pluripotent stem cells. Pluripotent stem cells are either embryonic stem cells extracted from embryos or induced pluripotent stem cells obtained by reprogramming adult stem cells. Organoids may develop both from normal cells and from tumor cells. In the latter case they are also called tumoroids. In order to check the effects of radiation on a given organ, it is preferable to use normal cells. In clinical applications, the mutations leading to cancer are introduced in organoids derived from normal tissue through applying CRISP/CAS9 mediated genetic editing or induction with the help of viruses (such as for instance the lentivirus). This allows reproduction of different stages in the genesis of cancer. In the present context, the source of genetic alterations is ionizing radiation of various types, especially HZE radiation the effects of which are still not very well understood. The idea of using organoids to investigate the stochastic effects due to ionizing radiation is still in its infancy, but it looks very promising. With organoids it is in fact possible to mimic many human organs like the colon, pancreas, stomach, retina and many others. A more complete list of organs that have been simulated thus far can be found in [79]. It is worth noticing that the majority of techniques applied in clinical tests for detecting the genetic mutations and genomic instabilities mentioned in the previous Subsection can be applied to organoids also. Moreover, organoids have a long life (they can last for a few hundred days). This is very useful in verifying the effects of doses that are not acute as is the norm in BLEO. Of course, organoid technologies are still in development. As such, they have still some shortcomings that affect their application in medicine. The most important limitation in the present context is that organoids are not able to reproduce the full characteristics of in vivo organs. The original stem cells differentiate in organoids into different types of cells and many structures of the real organ are reproduced, although not all of them. Recently, the induction of myelinating oligodendrocytes in cortical spheroids recapitulating the human brain has been possible [80]. In this situation it is not clear if it will be possible to study all consensus molecular subtypes CMS1-4 of CRC mentioned in the previous Subsection because a particular subtype could originate from a kind of cells that is lacking in organoids. In principle, it could be possible to stimulate the growth of organoids with that particular kind of cell. In practice, however, despite the dramatic improvements in our understanding of CRC thanks to molecular medicine, it is not yet recognized which kinds of cell tumors of a single molecular subtype are starting. On the positive side, organoids are based on human cells. For this reason, they are certainly suitable to study tumorigenesis following exposure to ionizing radiation in humans. Animals, even humanized mice, have not the same genetic characteristics as humans, so they also have a different oncogenic process.

## Conclusions

The probabilities of the occurrence of deterministic and stochastic effects during a typical mission to the Moon or Mars have been estimated using statistical and physical arguments. The probability of deterministic effects turns out to be very low. On the contrary, the ERR of inducing fatal cancer due to the radiation in space is high (ERR_space_∼1.75 Sv^-1^). The advantages and limits of these results has been discussed. In particular, it has been stressed that the formulae for estimating deterministic effects are not suitable for non-acute doses of HZE particles. The effects due to these particles are quite difficult to estimate due to the lack of statistics. A new strategy has been proposed that exploits 3D models and the recent advances of molecular medicine to detect the early stages of cancer formation in human cells. 3D models are quite efficient in capturing the characteristics of human organs, while the methods of molecular medicine have the potential to shorten the time necessary to predict the risks of cancer after irradiation. The initial steps of the development of cancer can be monitored looking at a limited set of genes that are mutated or epigenetically silenced together with the appearance of genomic instabilities like CIN or MSI. This increases the chances to fill, to some extent, the gap in our knowledge about the processes behind the onset of cancer, starting from the physical and chemical effects visible immediately after exposure to radiation, like double strand breaks, oxidation of nucleotides due to ROS etc. and the later appearance of stochastic effects like solid tumors. As an application of the proposed strategy, we find a deeper interpretation of the mechanisms that cause oxidative stresses in cells linking them with the activation of the WNT pathway in cells, a known signaling pathway that becomes upregulated in the earlier stages of colorectal cancer.

## Acknowledgements

The authors wish to thank heartily B. Górski and S. McKenna-Lawlor for fruitful discussions. The research of FF has been supported by the Polish National Science Centre grant 2020/37/B/ST3/01471. FF would also like to acknowledge the contribution of COST Action CA17139.

## Footnotes

1 In [34] a late effect of ionizing radiation like the early appearance of cataracts has been studied. From the statistics obtained examining 295 astronauts involved in several different missions it turns out that radiation increases the risks for cataracts in a dose and LET dependent way.

2 In [42] a low dose is defined as any dose from background to 150mSv. Other definitions of low doses based on different criteria and methodologies can be found in [40] – Annex G.

3 This value is the result of the sex-averaged estimates of the ERR at age 70 after exposure at age 30 given in Ref. [49].

4 Actually, while the Hiroshima bomb was plutonium based, the Nagasaki one was uranium based and its percentage of neutrons was less by at least by a factor 2 ([49]).

5 In vitro experiments are easy to perform, but have strong limitations. The cells used in such experiments are very different from those of the human body: They duplicate in a different way, the cell polarity is lost, the cell differentiation that distinguishes the cells in an organ is lost just to mention a few disadvantages.

6 Some genes involved in signal transduction pathways play also a role as oncogenes or historically have been classified as oncogenes. In consequence, sometimes genes such as PIK3CA, which belong to the PI3K signaling pathway, are referred to in the literature as oncogenes.

7 Telomeres are structures consisting of a repetition of the DNA sequence TTAGGG. They are found at both ends of each chromosome and protect chromosomes from nucleolytic degradation, from inappropriate recombination, repair and fusion. The role played by the shortening of telomeres in making the genome unstable has been discussed in [14]. It is noted that exposure to radiation may also activate telomerase in human somatic cells stimulating the increase of the length of telomeres. The lengthening of telomeres has indeed been observed in irradiated limphoblasts [69] and, more recently, also in humans, see the NASA Twin Experiment reported in [70].

8 More information about the transforming growth factor TGF-ß in CRC can be found for instance in [72].

9 Hypermutation means that cells have a heavy burden of mutations. In CRC this means a mutation number equal or greater than 12 mutations/Mb [76], where Mb denotes mega base pairs. Usually tumors with MSI are hypermutated because of the errors in MMR, while chromosomally unstable tumors (CIN dominated) are not hypermutated.

10 Upregulation of a gene means that the expression of that gene is increased, leading to an increase of the amount of its products. In signal transducing pathways several genes are involved.

11 Organoids are not to be confused with other 3D cell aggregates like for instance spheroids, which are not able to mimic the structure of in vivo organs [79].

